# Functional diversity in the output of the primate retina

**DOI:** 10.1101/2024.10.31.621339

**Authors:** A. Kling, S. Cooler, M.B. Manookin, C. Rhoades, N. Brackbill, G. Field, F. Rieke, A. Sher, A. Litke, E.J. Chichilnisky

## Abstract

The visual image transmitted by the retina to the brain has long been understood in terms of spatial filtering by the center-surround receptive fields of retinal ganglion cells (RGCs). Recently, this textbook view has been challenged by the stunning functional diversity and specificity observed in ∼40 distinct RGC types in the mouse retina. However, it is unclear whether the ∼20 morphologically and molecularly identified RGC types in primates exhibit similar functional diversity, or instead exhibit center-surround organization at different spatial scales. Here, we reveal striking and surprising functional diversity in macaque and human RGC types using large-scale multi-electrode recordings from isolated macaque and human retinas. In addition to the five well-known primate RGC types, 18-27 types were distinguished by their functional properties, likely revealing several previously unknown types. Surprisingly, many of these cell types exhibited striking non-classical receptive field structure, including irregular spatial and chromatic properties not previously reported in any species. Qualitatively similar results were observed in recordings from the human retina. The receptive fields of less-understood RGC types formed uniform mosaics covering visual space, confirming their classification, and the morphological counterparts of two types were established using single-cell recording. The striking receptive field diversity was paralleled by distinctive responses to natural movies and complexity of visual computation. These findings suggest that diverse RGC types, rather than merely filtering the scene at different spatial scales, instead play specialized roles in human vision.

## Introduction

The functional diversity of the many visual pathways emanating from the retina, and their distinct roles in vision, remain poorly understood. In mice, diverse retinal circuits mediate selective responses to ecologically relevant stimuli, including looming motion and object/background segregation^1–5^. These and other specialized signals are conveyed to numerous targets in the brain by over 40 distinct retinal ganglion cell (RGC) types^4,6^. In primates, much less is known. The four numerically dominant primate RGC types exhibit textbook center-surround spatial filtering properties^1–5,7^, suggesting that more complex aspects of visual processing may occur primarily in the brain^8–10^. Yet it is unclear whether the remainder of the ∼20 molecularly and anatomically distinct primate RGC types perform similar functions or instead exhibit non-classical properties^11–17^. Recent work on a few primate RGC types has suggested that the primate retina may transmit more complex visual signals than previously appreciated^14,18–21^. However, the majority of primate RGC types have not been functionally characterized. Thus, despite decades of research on the parallel visual pathways in primates, the functional diversity of the RGC types that compose them remains largely unknown.

## Results

To probe the functional diversity of retinal ganglion cell (RGC) types in primates, we performed large-scale data mining on a collection of nearly one thousand 512-electrode recordings from macaque retina *ex vivo*, a unique database capturing responses from ∼500,000 RGCs, as well as twelve recordings from the human retina. This effort benefited from novel spike sorting tools^22^, and the results were confirmed using the analysis below. Most of the recordings revealed a variety of distinct RGC types in a single patch of the retina (Fig. 1, Suppl. Figs. 1,2). These cell types were identified by stimulating the retina with spatiotemporal noise and either manually grouping or quantitatively clustering cells based on three aspects of the recorded responses: (1) the spike-triggered average (STA) stimulus (see Methods), a convenient summary of the spatial, temporal and chromatic aspects of light response^23,24^, the electrical image (the spatiotemporal voltage fingerprint of spikes produced by each cell), and (3) the interspike interval distribution, which captures the temporal structure of spike trains that help to distinguish cell types^25^.

**Figure 1.**
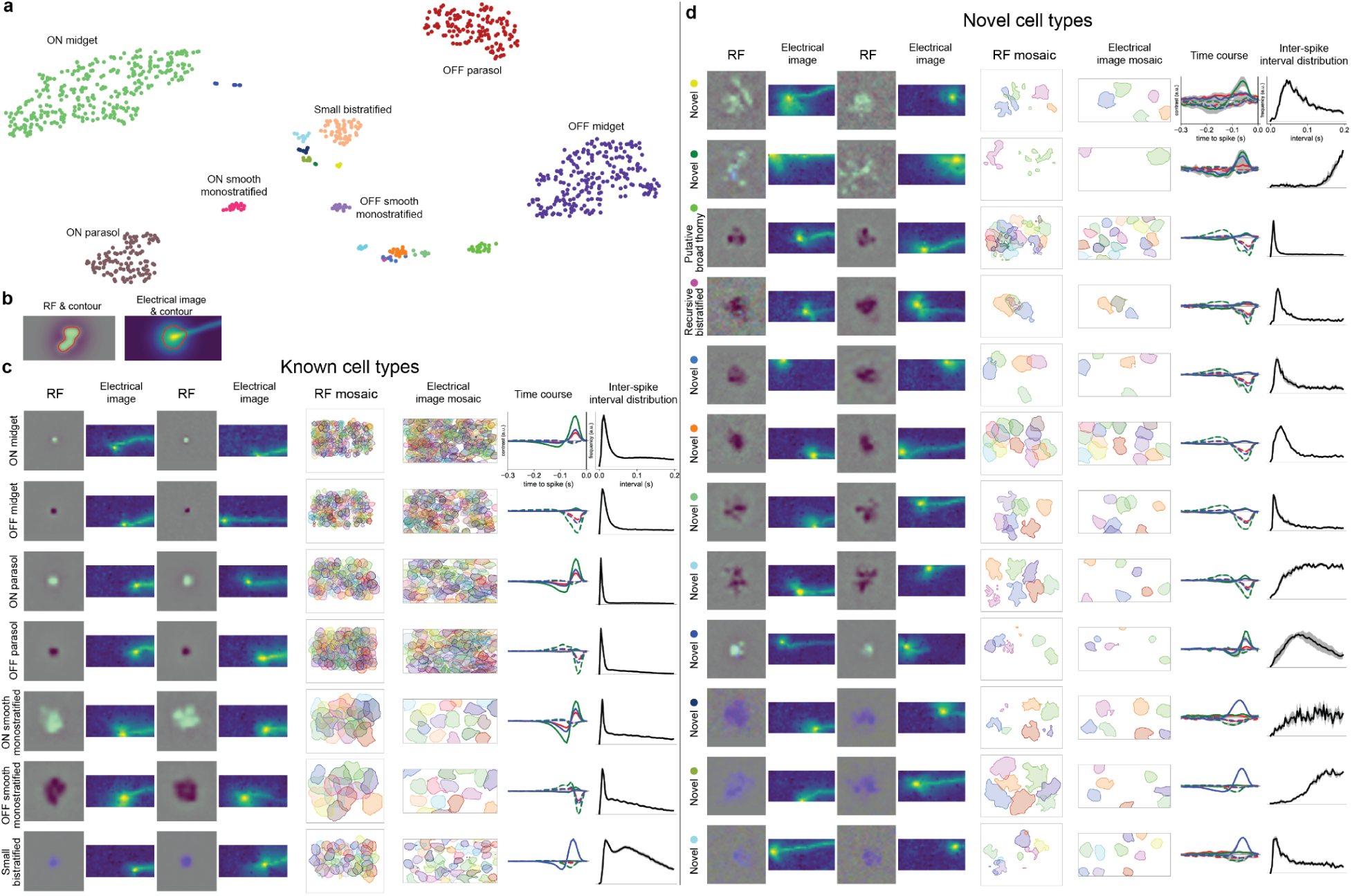
Identification of novel RGC types from a single recording (9.5 mm equivalent temporal eccentricity) in macaque retina. **a.** Representation of intrinsic (interspike interval distribution) and light response (time course) parameters (see text) of 938 simultaneously-recorded cells, represented in two dimensions using t-SNE. Cells fall into distinct clusters which are then further divided into distinct putative cell types by manual examination (see Methods). **b.** Illustration of RF and electrical image contour generation. **c.** Functional properties of cell types known from previous studies. For each cell type, panels from left to right show spatial RF and electrical image for two example cells, and RF contours, electrical image contours (somato-dendritic component), time course, and interspike interval distribution for all cells of this type. For time course and interspike interval distribution plots, mean across cells (lines) and standard deviation (gray shading) are shown. Time course includes an average of red, green, and blue display primaries in the STA pixels with the positive (ON time course, solid lines) or negative (OFF time course, dashed lines) peak preceding the spike (see Methods). RF images are 1200 µm wide. **d.** Same as **c**, for 12 putative novel cell types identified manually.

**Figure 2.**
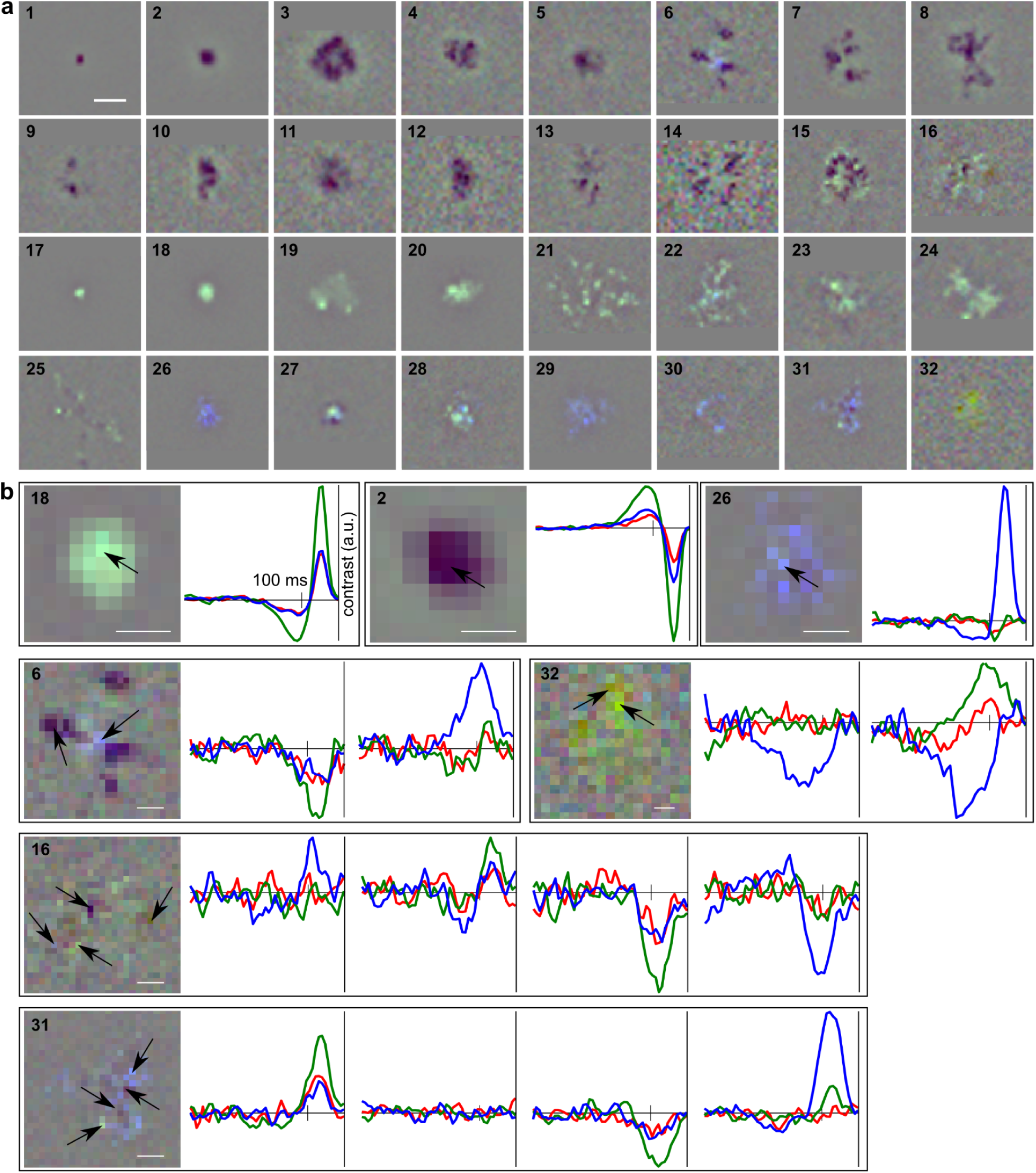
Diversity of spatial RF structure of primate RGCs, including known and novel types. **a.** STAs of 32 representative cells of distinct putative types from three recordings. Panels with identified cells: 1 - OFF midget; 2 - OFF parasol; 3 - OFF smooth monostratified; 4 - putative broad thorny (see Fig. 4); 5 - recursive bistratified (see Fig. 4); 17 - ON midget; 18 - ON parasol; 19 - ON smooth monostratified; 26 - small bistratified. Scale bar: 500 µm. All cells except one (panel 32) were selected from two recordings. The equivalent temporal eccentricity of the three preparations was 8mm, 10mm, and 8mm; the RF diameters of OFF parasol cells were ∼300 µm in all three recordings. If the RF was close to the edge of the visual stimulus field, a gray background around the edges was added for illustration purposes. **b.** Time courses of cells selected from **a** (panel numbering corresponds to **a**) computed from single STA pixels (arrows). ON and OFF parasol cells (panels 18 and 2) show time courses of the red, green and blue display primaries with relative amplitudes corresponding to L/M-cone inputs, while the small bistratified cell (panel 26) shows red, green and blue time course amplitudes corresponding to their known dominant S-cone input. Panels 6, 16, 31, and 32 demonstrate a variety of spectral inputs, polarity, and kinetics in different parts of the RFs of novel cell types (panels correspond to arrows in left-to-right order), indicating various mixtures of L/M and S-cone signals at various locations in their RFs. Scale bars: 200 µm.

The cells in each functionally-defined cluster exhibited highly stereotyped properties. Some clusters, corresponding to the seven RGC types previously identified in MEA recordings (ON and OFF midget cells, ON and OFF parasol cells, small bistratified cells, and ON and OFF smooth monostratified cells^14,19,24,26–28^, were easily identified (Fig. 1c, Suppl. Figs. 1,2). Together these cell types comprise over 70% of all primate RGCs^26–28^. As observed previously and as expected from anatomical work, the spatial receptive fields (RFs) of these cells tiled the retina with minimal overlap^14,19,24–26,29–32^. However, in most recordings, additional functional clusters were also observed (Fig. 1a, Suppl. Figs. 1,2). Cells in most of these clusters exhibited properties that distinguished them clearly from the aforementioned seven RGC types, and their RFs and somato-dendritic parts of their electrical images also tiled the retina (Fig 1d). Many of these clusters likely correspond to known morphological and/or molecularly identified types, but for most the correspondence is unknown (see below). Therefore we will refer to the additional clusters as (functionally) “novel” types.

The novel RGC types exhibited a surprising and striking variety of spatial RF properties (Fig. 2). As expected based on anatomical data, RF sizes varied greatly, with the largest RFs ∼20x larger in diameter than midget cell RFs. Also as expected, parasol cell and midget cell RFs exhibited smooth, symmetric, center-surround structure (Fig. 2a, panels 1-2, 17-18), in agreement with the textbook view of spatial filtering by RGCs^2,7^. However, the RFs of simultaneously-recorded novel types exhibited a range of unexpected properties. All novel cell types exhibited asymmetric and irregular RF outlines, sometimes including branching organization, potentially reflecting the structure of their dendritic arbors (Fig. 2a, panels 6-8, 13-14, 25)^14,33^. Most cell types exhibited a spatially patchy RF, with multiple islands of high light sensitivity separated by gaps of low light sensitivity, a property previously seen in only two macaque cell types (ON and OFF smooth monostratified cells - Fig. 2a, panels 3,19)^19^. These patches were more sparsely spaced in some cell types than others (Fig. 2a, compare panels 6-7, 13-14, 21-22, 25, 30). In some cell types, the patchy structure consisted of spatially segregated ON (bright) regions and OFF (dark) regions, arranged in a manner inconsistent with center-surround organization (Fig. 2a, panels 6, 15-16, 25, 27, 31). Importantly, these unusual features were highly stereotyped in cells of each type, and very different in cells of different types. They were also not visible in the RFs of simultaneously-recorded parasol cells covering the same region of the retina recorded (not shown), as seen in previous work^19^, indicating that they are not attributable to photoreceptor damage. Although only 6 human retinas were examined, qualitative inspection suggests that they exhibit diverse RF structures similar to those in macaque (Suppl. Fig. 2).

The chromatic properties of novel RGC types were also striking and surprising. The strength of short-wavelength sensitive (S) cone inputs compared to long- and middle-wavelength sensitive (L/M) cone inputs was assessed by the relative magnitude of the three color primaries in the STA (see Methods; Fig. 2b, panels 2, 18, 26). S-cone inputs varied widely across cell types, and varied across the RF in certain cell types (Fig. 2b, panels 6, 16, 31, 32). Specifically, six RGC types exhibited strong/predominant S-ON input (Fig. 2a, panels 26-31), two RGC types exhibit strong S-OFF input (Fig. 2a, panels 16, 32), and nine types received weaker/occasional S-ON or S-OFF input – many more than the five RGC types previously known to receive measurable S-cone input^34–37^. The eight cell types with strong/predominant S-cone input also frequently received L/M-cone input, of ON and/or OFF polarity, either co-localized with or spatially segregated from the S-cone inputs (Fig. 2b, panels 16, 31, 32). In one cell type, S-ON input was consistently localized in a small patch roughly in the middle of the RF, with strong L/M-OFF inputs forming surrounding branches or patches (Fig. 2a and b, panel 6). Interestingly, S-OFF inputs were less common than S-ON inputs, and were usually located in small, weaker patches within larger, L/M-cone dominated RFs. Only two very rarely recorded low-density cell types exhibited dominant S-OFF input throughout the RF (Fig. 2a, panel 32) or large patches with S-OFF input (Fig. 2a, panel 16). The cells of the first type also exhibited L/M-ON patches within the RF (Fig. 2b, panel 32), while cells of the second type received S-ON, L/M-ON, and L/M-OFF inputs (Fig. 2b, panel 16). Strong S-OFF and L/M-ON inputs in these two cell types make them possible candidates for the previously described inner and outer intrinsically photosensitive cells^35^; their rare appearance in the recordings matches very low reported density (3-5 cells/mm^2^ combined). Together, these results indicate that the complexity of S-cone signal processing in the primate retina is much greater than previously appreciated.

The striking RF spatial structure observed in the novel primate RGC types could potentially reveal specific functions in natural vision, perhaps analogous to the cell type specific selectivity for ecologically relevant stimuli seen in other mammalian retinas^3,5,38^. As a first test of this possibility, the firing patterns of novel RGC types were examined in response to a sequence of naturalistic images jittered according to the statistics of fixational eye movements. Multiple distinct response patterns were observed: some cells fired at a high rate continuously during image jitter, while others responded preferentially to image transitions (Fig. 3a,b). Details such as response onset, duration, spike count, and peak firing rate averaged across all presented images were highly conserved within each cell type but differed substantially across cell types, further corroborating cell type classification (Fig. 3a,b). In addition to these features of the average response, the different RGC types varied greatly in the reliability of their responses to repeated stimulation with the same image (Fig. 3c, abscissa). To probe nonlinearity of light responses in each cell, the linear projection of the visual stimulus image against the STA was computed for all images, and the evoked responses were ranked according to this linear prediction. RGCs of multiple cell types responded strongly to images with most negative linear predictions (Fig. 3a, bottom of each raster), revealing nonlinear light response characteristics that are not seen in better-known cell types such as parasol cells (Fig. 3a, left) (see ^18^). Thus, the diversity of RFs observed across the novel cell types is paralleled by diverse patterns of responses to complex images, suggesting the need for a deeper investigation of their contributions to natural vision^39^.

**Figure 3.**
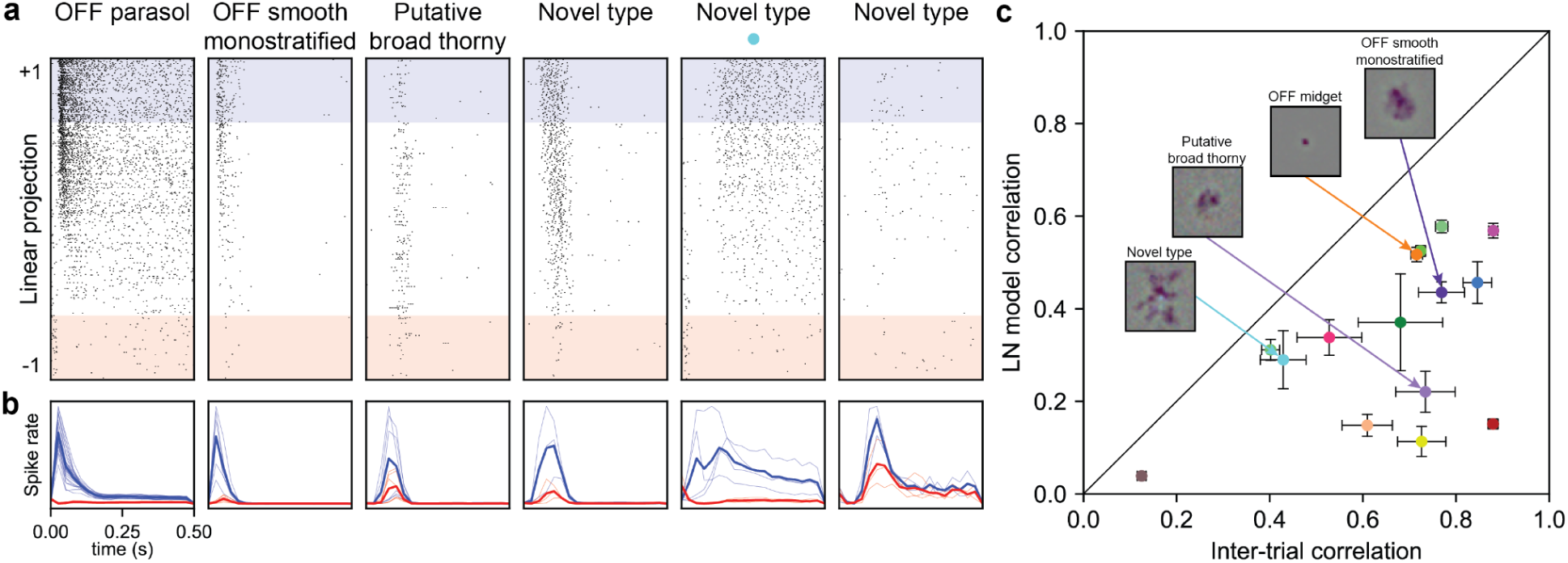
Diversity of responses to natural images across macaque RGC types. **a.** Rasters of responses to natural images for six example cells of different types, sorted in decreasing order by the linear projection of the stimulus onto the STA. Blue and red backgrounds denote 20% of the trials with the highest (blue, most positive) and lowest (red, most negative) values of the linear projection. **b.** Spike rate over time in responses to natural scenes, averaged across 20% of the trials with the highest (blue) and lowest (red) values of the linear projection. Thin lines: individual cells of the same type as in **a**; thick lines: average across all cells of this type. Time bins are 25 ms. **c.** LNP model performance as a function of trial-to-trial response reproducibility across cell types. The average correlation coefficient is shown for responses to repeated presentation of natural images with the average response (abscissa) or the response predicted by the LNP model (ordinate). Some cell types with very different RF sizes are similarly well-explained by the LNP model (e.g. OFF midget and OFF smooth monostratified cells), while others with similar RF sizes exhibit different model performance (e.g. OFF smooth monostratified and putative broad thorny cells (see below)). Points and error bars are mean ± standard error, *n* = 252 cells.

However, the observed computational complexity of light responses in different cell types was not easily predicted by their RF structure. For example, for some well-known and novel RGC types with very *different* RF size and spatial structure (parasol and midget vs. novel type denoted by cyan dot), responses were relatively well-explained by a simple pseudo-linear model: the trial-to-trial deviation of spiking responses from model fits was similar to the deviation from the average response (points closer to the diagonal in Fig. 3c). In contrast, for other well-known and novel RGC types with *similar* RF size and spatial structure, (OFF smooth monostratified vs. putative broad thorny cell (see below)), the performance of pseudo-linear models captured trial-to-trial variability effectively in some cell types (points closer to diagonal in Fig. 3c) but not in others (points far from diagonal). In summary, the sizes and complexity of RFs of each cell type did not reliably indicate the degree of response linearity. These findings motivate the need to establish the roles of mechanisms such as subunit nonlinearities or direction selectivity in creating the surprising responses of novel RGC types to natural images^40–43^.

Identification of the functionally novel cell types with known anatomically defined types could help elucidate the circuitry and mechanisms underlying complex RF structure and non-linear response properties. One such identification is the recursive bistratified cell. Based on previous work, recursive bistratified cells are known to be the only RGC type gap-junction coupled to A1 polyaxonal amacrine cells^13,14,18,44–48^. In single-cell recordings, this unique gap junction coupling was confirmed by intracellularly injecting anatomically identified recursive bistratified cells with fluorescent dye and observing several nearby labeled A1 cells (Fig. 4a; see ^18^), and by injecting A1 cells and observing coupling to recursive bistratified cells but not to other RGCs (not shown). In MEA experiments, only one novel RGC type displayed a distinctive two-peaked, short-latency cross-correlation of spike times with simultaneously recorded A1 cells^49^, a signature of electrical coupling^50,51^ (Fig. 4b). This signature was quantified using a two-peak cross-correlation metric (Fig. 4c), unambiguously identifying recursive bistratified cells in the MEA recordings (*n* = 164 cells, 28 recordings).

**Figure 4.**
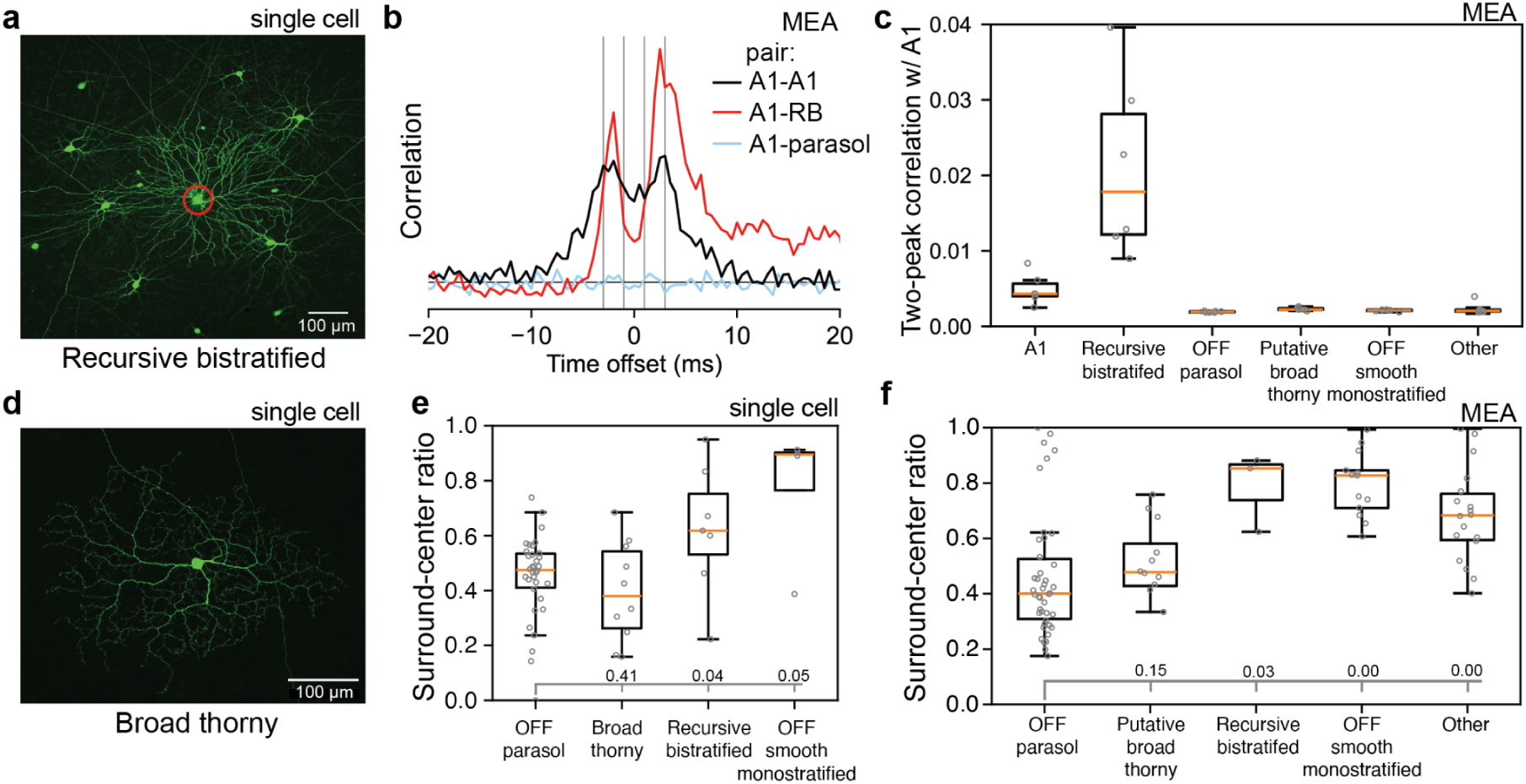
Comparison of broad thorny and recursive bistratified cell properties identified in single cell recordings and MEA recordings. **a.** Two-photon image of a dye-filled recursive bistratified cell from single cell recordings. Target cell soma is circled in red; other somata are A1 amacrine cells that are tracer-coupled by gap junctions. **b.** Pairwise cross-correlation histogram of spike times of an A1 amacrine cell and three other simultaneously-recorded cells: another A1 cell, a recursive bistratified cell, and an OFF parasol cell. Cross-correlations between A1-A1 and A1-recursive bistratified cells, but not A1-OFF parasol cells, exhibit two sharp peaks indicative of electrical coupling. **c.** Median nearest-neighbor (highest) two-peak cross-correlation with A1 amacrine cells, accumulated over six recordings, for OFF parasol, putative broad thorny, recursive bistratified, OFF smooth monostratified and other OFF-dominated cells (excluding midget cells) from MEA recordings. The correlation metric is defined as the lesser of the left and right peaks, minus the central trough. **d.** Two-photon image of a dye-filled broad thorny cell from single cell recordings. **e.** RF surround strength comparison in OFF parasol, broad thorny, recursive bistratified, and OFF smooth monostratified cells, measured in single cell recordings (*n* = 5). The surround-center ratio is computed as the ratio of the response to a full field stimulus to the maximum response to spots of varying sizes. Broad thorny cells are not significantly different from OFF parasol cells, unlike other cells tested (p values for each comparison are marked at the bottom of the plot) **f.** RF surround-center ratio comparison for OFF parasol, putative broad thorny, recursive bistratified, OFF smooth monostratified, and accumulated other OFF-dominated cells (excluding midget cells) from MEA recordings. The surround-center ratio is computed as the ratio of the summed contrast of the STA across space to the maximum summed contrast over circles of varying diameter. Similar to single-cell measurements, all cell types except putative broad thorny cells are significantly different from OFF parasol cells (p values for each comparison are marked at the bottom of the plot). Each data point is the median value for all cells of that type within a single recording (*n* = 42 recordings). In all box and whisker plots the upper and lower lines of the box are the first quartile and the third quartile of the data, with an orange line at the median. The whiskers are drawn at the farthest data point lying within 1.5 times the inter-quartile range from the box. Data points are drawn in gray circles.

A second novel RGC type, the broad thorny cell, was tentatively identified based on several lines of evidence^13,14,18,20,44–48^. In single-cell recordings, the anatomically identified broad thorny cells (Fig. 4d) exhibited RF surround suppression (measured using responses to spots of varying diameter) approximately as strong as that seen in OFF parasol cells (Fig. 4e)^18^ and substantially stronger than that seen in OFF smooth monostratified and recursive bistratified cells (Fig. 4e) and other sampled cells (not shown). The broad thorny cells also had substantially slower responses than OFF parasol or recursive bistratified cells. In MEA recordings, several OFF-dominated cell types had RF sizes and spatial structure, slow and biphasic STA time courses (Fig. 1), and ON-OFF response characteristics (Fig. 3b) consistent with properties of broad thorny cells in single cell recordings (see ^18^). However, only one of these types had RF surround suppression (measured using the STA integrated over increasing diameters) comparable to that of OFF parasol cells (Fig. 4f). Additionally, the putative broad thorny cells, like the recursive bistratified and ON and OFF smooth monostratified cells, tiled the region of recorded retina with their RFs and with the somato-dendritic components of their electrical images (Fig. 1d). While not definitive, this evidence supports the identification of broad thorny cells in MEA recordings (*n* = 389 cells, 33 recordings).

To understand the relationship between the functionally defined RGC types (Fig. 1) and previous anatomical surveys^11–14,28,52^, the total number of distinct cell types identified was estimated.

Although many novel cell types were readily defined by functional clusters of cells (Fig. 1a) with RFs and electrical images forming complete mosaics (Fig. 1d), other functional clusters contained cells with overlapping RFs and electrical images. Therefore, an algorithm was developed to estimate the number of distinct cell types in these mixed groups. The algorithm estimated the minimum number of cell types that could compose a group of cells without producing electrical image overlap within a type. The algorithm was calibrated using the electrical image overlap of identified low-density RGC types that tile the retina – ON and OFF smooth monostratified, recursive bistratified, and putative broad thorny cells (see Methods) – and was applied in two analyses below.

First, in each recording, the nine identified types – ON and OFF midget, ON and OFF parasol, ON and OFF smooth monostratified, small bistratified, broad thorny and recursive bistratified cells – as well as all cell types identified from functional clusters (Fig. 1a) with RFs and electrical images forming mosaics were set aside. The estimation approach was then applied separately to each remaining cluster of cells, resulting in a minimum count of 23-25 types per recording (*n* = 3).

Second, to account for the fact that different recordings typically exhibit a different collection of types, data from several macaque preparations from similar retinal eccentricities (confirmed by similar parasol cell RF size) were combined. First, the nine identified cell types were set aside, as above. Among the remaining RGCs in each recording, three cell groups were identified: ON-dominated cells, OFF-dominated cells, and cells with substantial S cone input. The minimum number of distinct types in each group in each recording was computed using the above method. Finally, it was assumed that the cell types comprising each of these groups are the same across recordings, and therefore that the number of cell types comprising each group is at least as large as the largest value obtained in any recording. This assumption is empirically supported for the well-known seven RGC types, though previous studies point to the possibility of exceptions^53,54^. Under this assumption, 23 distinct cell types in addition to the identified nine were observed, providing an estimate of 32 RGC types in the primate retina (Fig. 2).

## Discussion

The huge diversity of RF structure in novel RGC types (Fig. 2) is surprising, given the widely held view that the primary role of the primate retina in visual perception and behavior is bandpass spatial filtering produced by center-surround RFs^2,9^. This classical idea may be approximately applicable in the case of parasol and midget cells^55^. However, an indication of highly non-classical RFs in primate RGCs was seen recently in the island-like RFs of smooth monostratified cells^19,56^, adding to examples from previous work^35,57^. Unexpectedly, the present results show that complex RF structure of this kind appears to be the rule rather than the exception (Fig. 2). Note that unlike previous findings of fine structure in parasol and midget cell RFs at the ∼10 µm (2 min of visual angle) spatial scale of cone photoreceptors^27,31^, the patchy structure observed in the novel RGC types is coarse, with distinctive spatial structure at the ∼100 µm (∼20 min of visual angle) scale, often exceeding the size of midget and parasol cell RFs. Therefore, it is unlikely to be filtered out by the physiological optics of the eye^7^ and is likely to be relevant for natural vision.

The implications of the observed diverse RF structure for understanding retinal circuitry could be substantial. Some RGC types that exhibit spatially intermixed ON and OFF islands of light sensitivity may acquire this unusual structure by sampling inputs from both the inner and outer sublaminae of the inner plexiform layer. Although this might be expected of the four primate RGC types that have bistratified or broadly stratified dendritic arbors^14^, 6-8 additional cell types with mixed ON and OFF inputs appear in MEA recordings, some of which may correspond to other broad and diffuse cells described anatomically^13,58^ or to monostratified cells displaying ON/OFF properties^35^. Moreover, responses to naturalistic movies suggest that some cell types that have predominantly OFF (or ON) RFs measured using white noise may also receive co-localized ON (or OFF) input, further expanding the list of ON-OFF cell types unaccounted for by anatomical classifications. Certain other observations about RF structure are also surprising. The radially branching RFs seen in some cell types could point to a large impact of RGC dendritic morphology on the RF^59^. Also, S-cone inputs appear to be much more varied than was previously appreciated, with varying strength across cell types and in the RFs of individual cells. These findings suggest an unexpected sampling of inputs from S-cone bipolar cells vs. other bipolar cells, and are in line with recent reports of diverse S-cone pathways^34,37,60^ that apparently contradict the simple picture of two primary S-cone sensitive RGC types (large and small bistratified cells) receiving dominant S-cone excitatory input opposed to L/M-cone suppressive input^13,58^.

The lower bound of 32 RGC types is higher than ∼20 expected from individual anatomical and molecular surveys^13–15,28,47,61^. Several factors could explain this discrepancy. First, the collection of cell types identified in each of the preceding studies may not be the same; for example, each study may sample a subset of types and molecular methods may reveal different types than anatomical methods. Second, newer studies describe a few cell types not identified previously, suggesting that methodological advances over time provide a fuller view^17,52^. Finally, some surveys were based on stochastic labeling of a relatively small number of cells^47,52^. The present estimate of 32 types is likely a lower bound and could also be incomplete. First, the electrical image overlap analysis is prone to underestimation due to unrecorded cells (see Methods). Second, some cells were not included in the analysis because they were rarely observed or did not produce a measurable response to white noise stimulation.

Although the above findings point to much greater diversity in primate RGC light responses than has generally been appreciated, the relationship to the diversity seen in mice, rabbits, and other non-primate species is far from clear. Recent transcriptomic analysis has shown homology between primate cell types and a subset of mouse cell types, but their functional homology has not been established^62^. The recent discovery of direction-selective RGCs in primates illustrates that close homology can exist at the morphological, molecular, and functional level^21^. Indeed, the recursive bistratified cell type identified here is another candidate direction-selective type^14,20^.

However, several factors suggest that close homology may be the exception rather than the rule. First, molecular correspondence between RGCs in mice and primates is less clear than it is for other retinal cell classes^15^. Second, most numerous RGC types in primates (ON and OFF midget cells) and their genetic orthologs in mice (ON and OFF sustained alpha cells) exhibit substantial differences (e.g. relative RF size and numerical prevalence^62^). Third, the known number of morphologically and molecularly distinct types in primates (∼20), although potentially an underestimate, is far lower than that in mice (∼40), suggesting that the functional diversity in mice is greater^6,15^. Fourth, the species exhibit different visual behaviors and occupy different ecological niches, factors that likely exert different evolutionary pressures and could produce different visual computations in RGCs. Thus, a key next step is to probe more thoroughly the responses of the diverse novel RGC types to naturalistic stimuli^39^.

The present findings suggest avenues for future work in relation to human vision. Recent work has shown that the numerically dominant and well-studied cell types in macaques (midget, parasol, and small bistratified cells) have clear functional counterparts in the human retina^62–69^. While there is evidence for strong anatomical and molecular homology between less-studied RGC types across primate species^15,17,28^, it is unclear whether they are functionally identical between macaque and human. Probing this possible homology could advance our understanding of human vision and improve the function of retinal implants for vision restoration, which would likely benefit from mimicking the neural code of the retina with cell-type specificity^70^.

## Methods

### Tissue preparation

Eyes were obtained from macaque monkeys terminally anesthetized by other laboratories in the course of their experiments, in accordance with Institutional Animal Care and Use Committee requirements at their respective institutions. After enucleation, the eyes were hemisected and the vitreous was removed. In some cases, the eye was incubated in oxygenated Ames’ solution (Sigma, St. Louis, MO) at 33 C, pH 7.4 with added human plasmin (0.015 mg/ml, Fisher Scientific, Hampton, HN) for 20 minutes to assist with vitreous removal. Recordings were performed using small (2×3 mm) segments of retina with the retinal pigment epithelium and choroid still attached, situated 7-17 mm from the fovea (6-12 mm temporal equivalent eccentricity, or 29-56 degrees of visual angle)^55,71,72^. The choroid was then removed down to Bruch’s membrane to improve oxygenation.

Human eyes were obtained from brain-dead donors through Donor Network West (San Ramon, Ca). Human tissue preparation for recordings was then similar to that of macaque monkeys.

### Recordings statistics

Overall, more than 800 recordings from macaque and 12 recordings from human retina were considered. 33 recordings from 19 macaques and six recordings from five humans were analyzed in detail. Fig. 1 and Suppl. Figs. 1-2 show one recording each. Cells from three different recordings were used for Fig. 2. Eleven recordings were included for natural image analysis, with one of them used in Fig. 3. All 33 recordings were used in Fig. 4. 19 recordings were used in the electrical image overlap analysis.

### Multi-electrode array recordings and spike sorting

Custom multi-electrode arrays (MEAs) were used for recording. The MEAs had 512 electrodes with 60 µm pitch, in a 16×32 isosceles triangular arrangement over a rectangular region covering approximately 1×2 mm, or approximately 4×8 degrees of visual angle^73^. The retina was placed onto the MEA ganglion cell side down and held in place with a permeable membrane (Thomas Scientific, Swedesboro, NJ) attached to a titanium plug. The retina was continuously perfused with oxygenated Ames’s solution at 31-33 C during the recordings, which lasted up to 18 hours. Only the data collected while there were no changes in the recording quality were used for the analysis, and the majority of the data was obtained in the first 6 hours of the recordings. Voltage traces were band-pass filtered, amplified, and digitized at 20 kHz using custom electronics^73^. Spike sorting was performed with Kilosort2 software^22^. In a few recordings, YASS spike sorting software was also used; the results were largely in agreement with those obtained using Kilosort 2^22,74^. Post-spike sorting quality controls were performed for each recorded cell: 1) contamination - cells were excluded if one or more of the following was observed: refractory period violations within 1 ms; multiple axons in the electrical image (see below); receptive field (RF, see below) consisting of a mixture of RFs of other cells; 2) duplication - cells with similar electrical images were merged. Spike sorting was done separately for each ∼30-120 minute stimulus sequence. Cells were cross-identified across stimulus sequences by correlating their electrical images, described below.

### Single-cell recordings

For the patch clamp experiments, recordings were performed from a retinal pigment epithelium-attached retinal preparation at temporal equivalent eccentricities of 4-8 mm (18-36 degrees) in macaque retinas with conditions closely matched to MEA recordings. Visual stimuli were generated using the Stage software package (https://stage-vss.github.io) and displayed on a customized Lightcrafter 4500 projector^75^. The peak wavelengths of the LEDs in the projector were 405, 565, and 630 nm. Recordings were performed using borosilicate glass pipettes containing Ames’ medium during extracellular (loose patch) recordings. Cells were filled with biocytin (0.5%; EZ-Link hydrazide, ThermoFisher) during whole-cell, voltage-clamp recordings with an intracellular solution containing (in mM): 105 CsCH_3_SO_3_, 10 TEA-Cl, 20 HEPES, 10 EGTA, 2 QX-314, 5 Mg-ATP, and 0.5 Tris-GTP, pH ∼7.3 with CsOH, ∼280 mOsm^33^. The retina was superfused with 95% O_2_/5% CO_2_ bubbled Ames solution at 32-35 C. Cells were filled with biocytin (0.5%; EZ-Link, ThermoFisher) for later anatomical imaging. Following recording, tissue was fixed by immersion in a solution containing 4% PFA for 30-45 min and washed with 1X PBS. Tissue was counterstained with Streptavidin-conjugated Alexa 488 to visualize filled cells. Cell nuclei were stained with DAPI and starburst amacrine cells were stained with antibodies against choline acetyltransferase. These stains allowed for accurate identification of dendritic stratification in the inner nuclear layer^76^. Images of filled cells were acquired using a SP8 confocal microscope (Leica) and used for anatomical verification. Tissue quality was assessed by recording contrast responses from parasol ganglion cells neighboring the targeted broad thorny or recursive bistratified cell. In sensitive tissue, parasol ganglion cells were expected to increase their spike rate by at least 20 spikes s^−1^ to a 5% contrast spot presented over the RF at a photopic mean luminance^75^. Cells recorded in retinal regions that did not meet this selection criterion were excluded from the manuscript. Spike responses were recorded with binary white noise stimuli refreshing at 60 Hz (pixel size, 20-30 μm on a side) for at least 35 min, and spatiotemporal RFs were recovered by cross-correlating the stimulus sequence with the cell’s spike output.

### Electrical images and deduplication

The electrical image for each cell was computed by averaging short voltage traces (200 samples, 10 ms) centered on the time of each spike. Electrical images are composed of distinct components (soma, dendrites, axon) that are easily identifiable based on the voltage waveform shape and spatial configuration^24,32,73^. The number of axons visible in the electrical image and the spike propagation speed was used to classify cells as ganglion cells, polyaxonal amacrine cells, or contaminated cells.

Electrical images were also used to identify duplicate copies of a single cell produced in spike sorting as well as to associate recorded spikes with the same cell across several stimulus runs. To handle duplicate cells, spikes were pooled from cells with electrical images that had a high correlation coefficient. The correlation threshold for pooling was chosen to minimize the number of incorrectly pooled cells relative to manually supervised pooling using other properties, including precise axon paths, light response properties, and temporal firing patterns.

### Visual stimulation

A gamma-corrected cathode ray tube display (Sony Trinitron Multiscan E100; Sony, Tokyo, Japan), resolution 640×480 pixels, refreshing at 120 Hz, was used to deliver visual stimuli produced and controlled by custom MATLAB software. The image was optically reduced to 3.4 x 2.5 mm (5.3 µm pixels) and focused on the photoreceptor layer after passing through the mostly transparent electrode array and other retinal layers. The focal plane was refined by adjusting the focus to maximize the overall spiking activity during a fine-grained rapidly flickering checkerboard stimulus. The stimulus produced on average 800-2200, 800-2200, and 400-900 photoisomerizations per second for the L, M, and S cones respectively^19,27,77^.

Two visual stimulus types were used for the analysis. Several white noise stimuli were presented in each recording for 30 to 60 minutes each, for 2-8 hours in total. In each stimulus refresh, the contrast for each pixel on the display was randomly selected from a binary distribution (mean ± 96%), independently for each display primary. To maximize stimulation of different cells, several spatio-temporal resolutions were used (spatial: 21.2-106 µm pixels; temporal: 20-120 Hz refresh). In some cases, the stimulus was also randomly jittered on a 5.3 µm grid at each refresh.

Natural movies were constructed using images from the ImageNet database and presented as described previously^32,78,79^. Briefly, images were converted to grayscale values with a common mean and resampled to 320 x 160 pixels, for a final scale of 10.6 × 10.6 µm per pixel at the retina. Each image was displayed for 500 ms (60 monitor frames) and was jittered according to Brownian motion with a diffusion constant of 10 μm^2^/frame, selected to roughly match the amplitude of recorded fixational eye movements in humans and other primates^80–82^. Each natural movie stimulus was composed of interleaved blocks of unique images shown once, 5000 total, and 150 images (each repeated ten times) interleaved.

### Spike-Triggered Average

RFs were summarized using the spike-triggered average (STA) stimulus obtained with white noise stimulation. For each spatial and temporal resolution of the white noise stimulus used, reverse correlation was used to compute the STA^23^. Because different cell types respond most strongly to stimuli at different spatial scales, a *multi-resolution STA* was also computed by upsampling the STAs for each cell obtained at different spatial and temporal resolutions in time and space to the finest common resolution, weighing by the spike count, and summing. Because stimuli at lower resolutions are spatially and temporally correlated relative to the stimuli with finer resolutions, the resulting combined STA is not an unbiased estimate of the linear RF at the finest resolution. However, this approach provided a representation of spatial and temporal properties of the cell with a high signal-to-noise ratio, allowing for clearer distinction of cell types.

Importantly, the RF properties used to distinguish the cells were largely unaffected by combining the STAs (Fig. M1).

**Figure M1.**
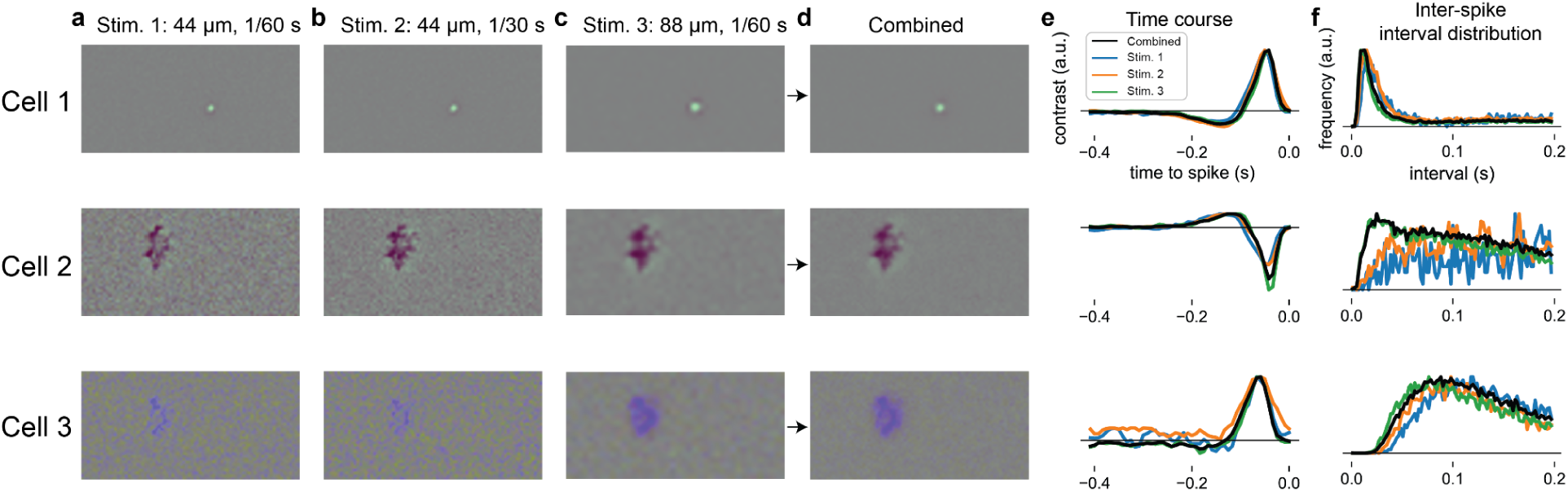
Multi-resolution STA improves signal-to-noise ratio but has no major effect on cell’s properties used for cell type identification. **a.** Spatial RF obtained from high spatial and low temporal resolution stimulus for three example cells. Full image width is 3.5 mm. **b.** Same, for the same spatial but higher temporal resolution stimulus. **c.** Same, for low spatial and high temporal resolution stimulus. **d.** Spatial RF obtained from multi-resolution STA (by combining single-resolution STAs in panels a-c). Note the noise reduction, but no major changes in the RF structure. **e.** Interspike interval distribution computed for single-resolution white noise stimuli (colored lines) and combined stimuli (thick black line), normalized. **f.** Largest time course obtained from single-resolution STAs (colored lines) and from combined STA (thick black line), normalized. Inter-spike interval distributions and time courses also do not exhibit major changes but the noise is reduced when combined STA is used.

### Cell type classification with intrinsic and light response properties

The spatial, temporal, and chromatic properties of light responses in each cell, as well as its intrinsic firing properties, were used in cell type classification.

The light response properties of each cell were measured from the multi-resolution STA (see above). The multi-resolution STAs were decomposed into spatial (RF) and temporal (time course) components. First, the red and green display primary components in the STA were averaged due to the significant overlap of L and M cone spectral sensitivity and the (presumed) indiscriminate sampling of L and M cones by all peripheral RGCs other than midget cells. An initial spatial RF was obtained by identifying pixels with contrast energy summed over the duration of the STA that deviated from the background noise energy distribution by more than a statistical threshold. Then, the spatial RF was subdivided into ON and OFF regions, separately for the RG color channel (sum of red and green display primaries) and the B color channel (blue primary), as follows. First, the time frame with the largest absolute contrast summed across all RF pixels was designated as the peak frame. Then, RF pixels with positive contrast at the peak time frame were identified as ON, and pixels with negative contrast as OFF, separately for the RG and B channels. Finally, time courses were computed separately for RG and B channels, and for ON and OFF pixels. For visualization of the RF, an inner product of the entire STA with the largest-amplitude time course was computed. For visualization of RF outlines and RF size computations, a contour was computed over the spatial RF, separately for each spatially continuous region of pixels statistically deviating from noise.

The intrinsic firing properties of each cell were summarized using the inter-spike interval distribution. A histogram of the time difference between each pair of adjacent spikes was created using 100 bins of 2 ms width.

### Data Selection

Some cells were eliminated from analysis because they exhibited properties strongly suggestive of failures of spike sorting. Some putative cells identified by spike sorting exhibited a refractory period violation, had a double axon in its electrical image, or exhibited RFs that could be decomposed into a linearly weighted sum of RFs of recorded midget and/or parasol cells. Such cells also often had inter-spike interval distributions and time courses resembling a mixture of midget and/or parasol cell spikes. Cells with any of these indications were omitted from analysis. Note that RFs of novel cells with hotspots of light sensitivity could not be decomposed to a linear sum of underlying midget or parasol RFs^19^, and these cells generally had inter-spike interval distribution and time course very different from those of midget and parasol cells. In addition, groups of very similar novel cells identified across the array exhibited similar inter-spike interval distributions and time courses. In some cases, the hotspots in the RFs of novel cells were smaller than the RF of midget cells, further indicating that they are unlikely to be a result of spike sorting error. Therefore these cells were included in the analysis.

### Clustering and Visualization

For visualization and clustering, parameters for each cell were obtained from the time course and RF. Time courses and inter-spike interval distributions for each recording were combined in a single vector for each cell and represented using 16 dimensions using PCA. The area of the convex hull of the RF of the color component with the highest STA amplitude was also used.

Nonlinear dimensionality reduction was applied to represent these 21-dimensional data in two dimensions (t-SNE and UMAP)^83–85^. Clustering was performed on this representation of the data using HDBSCAN and k-means^86^. Hyperparameters of dimensionality reduction and clustering were varied to identify groupings of cells that were stable across random initializations. For RF location visualization and mosaic display, green and red STA components were combined.

### Cross-correlation

Electrical coupling between cells was assessed from cross-correlogram of their spike trains, binned at 1 ms intervals. Previous work shows that electrically coupled cells display one or two narrow peaks in the cross-correlogram at 1-3 ms lag^50,51^. The following metric was used to compare broad stimulus-induced correlation with the precisely timed double peak signature in the cross-correlogram: the maximum value in the expected trough range ([-1; 1] ms) was subtracted from the smaller of the maximum values in the expected peak regions ([-3, -1] and [1, 3] ms). For each cell, this metric was calculated with all simultaneously recorded A1 cells, and the maximum value across A1 cells was used as a measure of the electrical coupling strength to the A1 network.

### Surround ratio

Two methods were used to estimate the center-surround ratio in the RFs of OFF-dominated cells. In single cell recordings, flashed negative contrast spots of multiple sizes centered on the RF were used. The surround ratio was calculated as the response to the full-field flash divided by the maximal response over all flashes. In MEA recordings, the spatial RF derived from the STA was summed within disks of variable sizes centered on the RF. The surround ratio was calculated as the full-field sum divided by the maximal sum over all sizes.

### Identification of S and L/M cone signals

To identify short-wavelength sensitive (S) cone inputs in the RFs of novel cells, a simple metric was devised based on the assumption that parasol cells receive only long- and middle-wavelength sensitive (L/M) cone inputs^27^. In the STAs of parasol cells, the maximal per-pixel amplitude of the blue display primary was 0.45±0.02 of the maximal amplitude of the green display primary. Hence, cells exhibiting an absolute value of the blue:green display primary ratio substantially greater than this value are likely to receive both S and L/M inputs in this pixel location. Indeed, small bistratified cells, which typically receive strong S-ON and weaker L/M-OFF input^26^, exhibited 2.19±0.3 (average per-pixel) blue:green ratio. A conservative threshold ratio of 0.75 was used to identify S-cone inputs.

### Manual cell type identification

Known cell types were identified first in each recording. ON and OFF midget, ON and OFF parasol cells, and small bistratified cells were identified based on their density, chromatic selectivity, and inter-spike interval distributions^26^. ON and OFF smooth monostratified cells were identified based on similarity of their time courses to those of parasol cells, their RF size and spatial structure, and their inter-spike interval distributions^19,24^. A1 amacrine cells were identified based on branching axons in their electrical images, RF size, compact RF spatial structure, and double peak in cross-correlation of their spike trains with each other^24,49^. Recursive bistratified cells were identified based on double peaks in cross-correlation of their spike trains with A1 amacrine cells. Broad thorny cells were identified based on their time course, RF size, center-surround ratio, and characteristic inter-spike interval distributions. Each of these cell types was verified by mosaic organization.

The remaining sparse cells were classified into putative types based on similarity of their time courses, inter-spike interval distributions, RF size and spatial structure. The similarity was assessed qualitatively by a human expert with over 10 years of experience analyzing MEA recordings from the macaque retina. This classification was conservative in the sense of attempting to minimize the number of putative distinct types. In many cases, the classification was straightforward and produced mosaics with minimal overlap. Most of these putative types could not be then combined with other groups without introducing mosaic violations. In other cases, cells in a single similarity group exhibited large, nonsystematic overlaps in electrical images and RFs. In these putatively multi-type groups, an electrical image overlap metric (see below) was used to determine the minimal number of cell types present in the group. Generally, it was possible to then manually identify non-overlapping putative types within these groups. A similar metric based on RFs was not used because their signal-to-noise ratio is generally lower than that of electrical images, and because the inhomogeneous structure of many RFs (Fig. 2) complicated the analysis.

### Cell type counting method

An analysis was developed to estimate the minimum number of putative cell types in a collection of recorded cells. The analysis was based on the assumption that electrical images of cells of each type do not overlap (see Fig. 1), and therefore that any given electrode can record a substantial signal from at most one cell of each type. Hence, the number of cells recorded on any given electrode provides a lower bound on the number of cell types. To implement this logic computationally, a cell was considered “present” on an electrode if (1) the electrode was less than 300 µm from the electrode with the largest signal amplitude recorded from that cell, and (2) the signal amplitude on the electrode was above a threshold fraction of the largest signal amplitude recorded from that cell on any electrode. The number of cells “present” on each electrode was computed, and the maximum of this value across electrodes was taken as a lower bound on the number of distinct cell types in the recording.The threshold used in the computation was adjusted such that, when this analysis was performed on a collection of low-density cells of four known types (ON and OFF smooth monostratified, recursive bistratified, and putative broad thorny), the analysis yielded approximately four types. Specifically, in 27 recordings with 7.3±2.1 cells of each of the four types per recording, a threshold fraction of 0.138 yielded an estimate of 3.96±0.8 types across recordings. This value of the threshold was then used to analyze other groups of recorded cells to determine a lower bound on the true number of cell types. This approach tends to underestimate the true number of cell types, because unrecorded cells can only reduce the estimate. For example, random subsampling of the four verified cell types to three or fewer cells per type per recording resulted in 2.4±0.5 identified types per recording. To verify that this counting method was not unduly affected by outlier electrodes, in the data sets and cell groups that yielded the largest cell type count, manual inspection was used to verify that the cells with overlapping electrical images were detected on multiple electrodes.

### Linear-nonlinear cascade model of light responses

To evaluate the linearity of RGC responses to natural images, a simple linear-nonlinear cascade model was fitted to experimental data^23,87^. First, the linear projection (inner product) of the spatiotemporal STA obtained from white noise (see above) and the training stimulus (5000 natural images, each shown for 60 monitor frames, ∼0.5 s) was computed at the monitor frame rate precision (8.3 ms). Firing rate during the stimulus was computed by convolving the spike train (at frame rate precision) with a gaussian kernel. Then, a static nonlinearity was computed by averaging the firing rate corresponding to the specific ranges of the linear projection values (the ranges were obtained by binning the linear projection such that each bin had the same number of instances). This nonlinearity was used to predict the firing rate in response to the testing stimulus (150 images, also shown for 0.5 s each). The resulting firing rate was correlated with the measured firing rate in response to each of the 10 repeats of the test images. The average correlation coefficient represented the performance of the model and was compared to the reliability of the measured response, given by the average correlation coefficient between the firing rate in each individual repeat and the average firing rate across all repeats.

### Statistics

For comparisons of surround-center ratio (Fig. 4) a two-tailed Mann–Whitney U test was used. Significance is marked in the panels.

### Software and Platforms

Data analysis was performed using MATLAB (The MathWorks, Inc.) and Python 3 with a set of additional modules: Numpy, SciPy, Scikit-Learn, Pandas, Matplotlib, UMAP, HDBSCAN, Jupyter Notebook^83–85^. Figures were arranged in Adobe Illustrator. Spike sorting was performed using Kilosort2^22^.

## Acknowledgements

This work was supported by NIH grants NEI R01-EY029247 to E.J.C., F.R., and M.B.M., R01-EY033870, R01-EY021271 and R01-EY032900 to E.J.C., Research to Prevent Blindness Stein Innovation Award to EJC; NEI R01-EY027323 to M.B.M., NEI R01-EY028542 to F.R., NEI P30-EY026877 core grant. We thank Eric Wu, Alex Gogliettino, and Jillian Desnoyer for technical assistance; T. Moore, S. Moriarty, the UC Davis Primate Center, and Washington National Primate Research Center for providing access to macaque retinas; William Harper and Donor Network West for providing access to human retinas; the Kilosort authors for their fundamental contributions to spike sorting technology.

## Author contributions

A.K. and E.J.C. conceived the work and designed the multi-electrode array experiments; A.K., C.R., N.B., and S.C. performed multi-electrode array experiments; M.B.M. and F.R. designed and performed single cell experiments; A.K., S.C. and E.J.C. analyzed the data; A.S. and A.L. provided multi-electrode array technology; A.K. and E.J.C. wrote the first version of the manuscript; all authors contributed to editing and producing the final version of the manuscript.

## Materials & Correspondence

Alexandra Kling

