## Supplemental Figures for "Functional diversity in the output of the primate retina"

### Supplemental Information

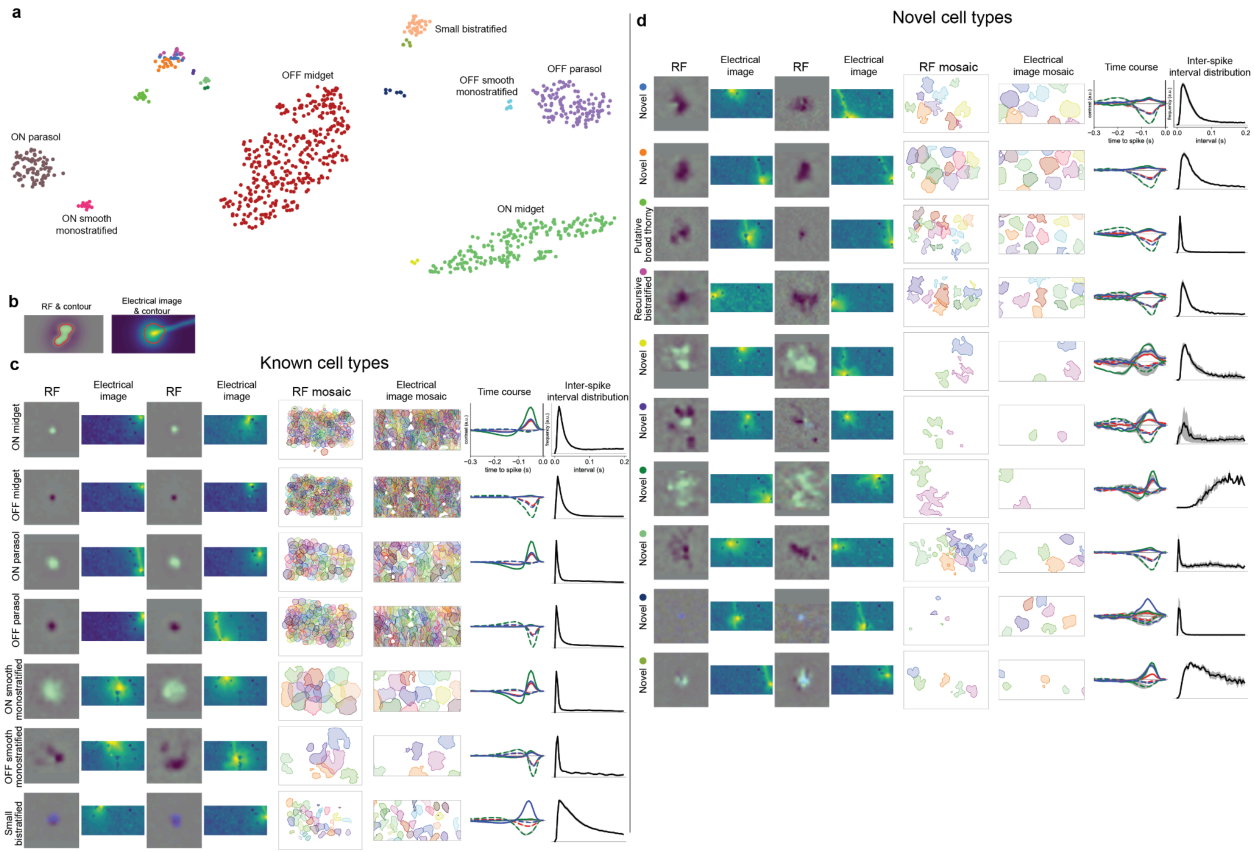


**Figure Supplement 1**. Identification of novel RGC types from a single recording (16 mm equivalent temporal eccentricity) in macaque retina. **a**. Representation of intrinsic (interspike interval distribution) and light response (time course) parameters (see text) of 914 simultaneously-recorded cells, represented in two dimensions using t-SNE. Cells fall into distinct clusters which are then further divided into distinct putative cell types by manual examination (see Methods). **b.** Illustration of RF and electrical image contour generation. **c**. Functional properties of cell types known from previous studies. For each cell type, panels from left to right show spatial RF and electrical image for two example cells, and RF contours, electrical image contours (somato-dendritic component), time course, and interspike interval distribution for all cells of this type. For time course and interspike interval distribution plots, mean across cells (lines) and standard deviation (gray shading) are shown. Time course includes an average of red, green, and blue display primaries in the STA pixels with the positive (ON time course, solid lines) or negative (OFF time course, dashed lines) peak preceding the spike (see Methods). RF images are 1200 µm wide. **d.** Same as **c**, for 10 putative novel cell types identified manually.

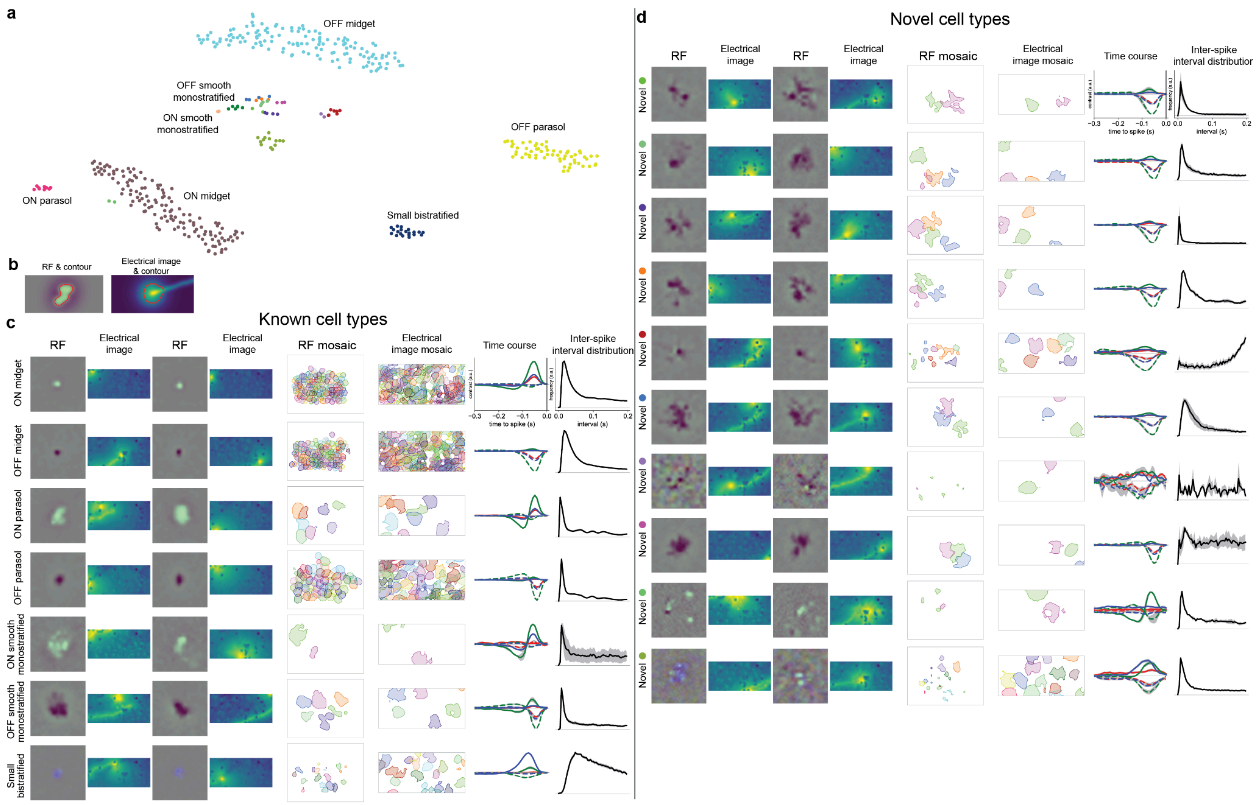


**Figure Supplement 2**. Identification of novel RGC types from a single recording (12 mm equivalent temporal eccentricity) in human retina. **a.** Representation of intrinsic (interspike interval distribution) and light response (time course) parameters (see text) of 410 simultaneously-recorded cells, represented in two dimensions using t-SNE. Cells fall into distinct clusters which are then further divided into distinct putative cell types by manual examination (see Methods). **b.** Illustration of RF and electrical image contour generation. **c.** Functional properties of cell types known from previous studies. For each cell type, panels from left to right show spatial RF and electrical image for two example cells, and RF contours, electrical image contours (somato-dendritic component), time course, and interspike interval distribution for all cells of this type. For time course and interspike interval distribution plots, mean across cells (lines) and standard deviation (gray shading) are shown. Time course includes an average of red, green, and blue display primaries in the STA pixels with the positive (ON time course, solid lines) or negative (OFF time course, dashed lines) peak preceding the spike (see Methods). RF images are 1330 µm wide. **d.** Same as **c**, for 10 putative novel cell types identified manually.
